# How to navigate the myosin-V motor through the actin network

**DOI:** 10.1101/2020.03.11.987495

**Authors:** Peter Spieler, Angela Oberhofer, Dennis Zimmermann, Edwin Brames, Alistair N. Hume, Zeynep Ökten

## Abstract

Myosin-V (MyoV) is a ubiquitous motor protein that transports an astonishingly diverse set of cargos on the actin network in eukaryotes. Phosphorylation-dependent processes often regulate MyoV-mediated cargo transport, molecular details of which remain largely unknown. We previously showed that phosphorylation regulates MyoV’s switching from microtubules onto actin filaments, not its motor activity. Regulation of switching at reconstituted microtubule-actin-crossings in fact sufficed to recapitulate the MyoVa-driven redistribution of pigment-organelles in amphibian melanophores. However, in those cells, MyoVa also encounter many actin-actin crossings. Here, we show that isolated MyoVa motors switch with equal probabilities at reconstituted actin-actin-crossings. Under the control of its adaptor-protein melanophilin (Mlph), however, the motor differentiates between the actin filaments at crossing points in a phosphorylation-regulated manner. Whereas phosphorylation of Mlph forced about ∼2/3 of MyoVa to ignore the intersections, dephosphorylation completely reversed this behavior and forced ∼2/3 to switch. We show that the filament-binding domain (FBD) of Mlph controls this switching behavior. This property evolved in amphibians, but not in the early vertebrate zebrafish. By protein engineering, we demonstrate that changes of a few residues are sufficient to impart actin-binding capability onto the zebrafish Mlph. We thus unmask the molecular beginnings of dual filament binding in Mlph that allow it to control the switching behavior of MyoVa at cytoskeletal crossings. We therefore propose a direct link between intracellular phosphorylation activity and the adaptor-protein, not to regulate MyoVa activity, but to navigate the motor through the entire cytoskeletal maze for correct positioning of cargo.

**Significance statement:** In virtually all eukaryotic cells, numerous myosin motors have to navigate through an elaborate actin network for timely transport of intracellular cargo. Here, we unmask an unintuitive regulation of the myosin-Va motor that is involved in pigment organelle transport. We demonstrate that myosin-Va differentiates between the same actin filaments and displays regulated switching at reconstituted actin-actin crossings, an unexpected behavior that has been predicted from previous theoretical work. We trace this regulation back to the adaptor protein of the myosin-Va motor and show that this regulation was present in amphibian but had not evolved in the early vertebrate zebrafish. Notably, we demonstrate that the evolution of actin-binding capability is achieved by changing a few residues in the adaptor protein.

## Introduction

How cells regulate kinesin, dynein, and myosin motors to ensure timely positioning of intracellular cargo at the right place is of outstanding interest. Instead of working alone, different motor types often team up on a given cargo. Particularly convoluted cases involve kinesin, dynein, and myosin motors simultaneously, all of which appear to know how to take turns to achieve a correct and timely positioning of cargo on the microtubule and actin cytoskeleton.

A well-known example of such convoluted transport occurs in melanophores from early and lower vertebrates to give rise to physiological color change. These specialized cells produce large, pigment-filled organelles, termed melanosomes, onto which they simultaneously recruit kinesin, dynein, and myosin motors (1-7). Fish and amphibians make use of the prototypical cAMP-PKA-dependent regulatory pathway to consciously control the deployment of these three different types of motors (Fig. 1A and B) (8-11). When external cues lead to an increase of intracellular PKA activity, melanosomes disperse throughout the cell on the actin network (Fig. 1A). Conversely, when PKA activity is decreased, and phosphatase activity increased, melanosomes switch from the actin onto the microtubule network to cluster in the cell center. This dynamic redistribution of large black granules in turn controls the light-scattering properties of the cell, allowing the organism to appear light or dark depending on external cues.

**Fig. 1:**
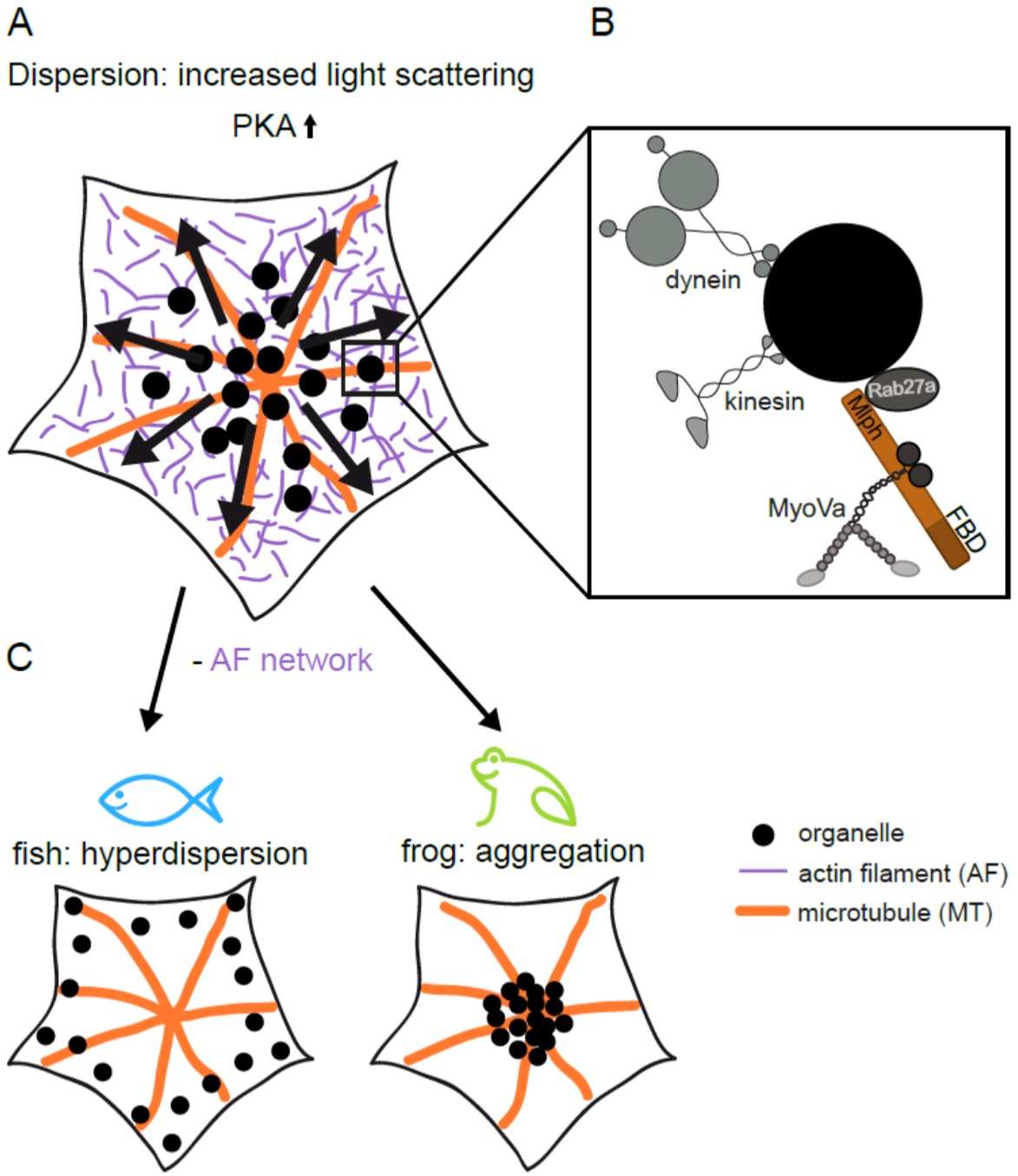
Fish and amphibians differ in use of the microtubule and actin networks in melanosome transport. (A) Illustration of melanosome dispersion in melanophores. The cell increases the scattering of light by dispersing the large black melanosomes as a response to increased PKA activity (black arrows indicate the overall direction of transport). (B) Kinesin, dynein, and myosin motors are found simultaneously on the melanosome surface. Recruitment of MyoVa is achieved by the Rab27a GTPase and the melanophilin (Mlph) adaptor protein (FBD indicates the C-terminal filament-binding domain). (C, left panel) Removal of the actin network prompts hyperdispersion in fish, i.e. the kinesins motor transport the melanosomes to the end of the microtubules. (Right panel) Frog fails to display hyperdispersion and melanosomes are transported to the cell center by the dynein motors after removal of the actin network, demonstrating the essential role of the actin network system to disperse melanosomes in amphibians.

However, while fish and frog models make use of kinesin, dynein, and myosin for regulated melanosome redistribution, they differ substantially in the functional use of the microtubule and actin cytoskeletal systems to achieve voluntary color change. Fish and amphibians both depend on the dynein motor to cluster their melanosomes in the cell center upon decrease of PKA activity (2, 3). Dispersal of the organelles as a response to increased PKA activity, on the other hand, requires the presence of the MyoVa/actin system in the amphibian (5), but not in fish model (12). In the absence of the actin network, fish melanophores efficiently move melanosomes outwards in a kinesin-depended manner on the microtubules upon increase of PKA activity (12). Fish is thus capable of the so-called hyperdispersion on the microtubule network, i.e. melanosomes are transported out towards the plus-ends of the microtubules (Fig. 1C, left panel). In a curious twist, removal of the actin network system in amphibian melanophores irreversibly clusters the melanosomes in the cell center in a dynein-dependent manner (Fig. 1C, right panel) (5), suggesting that the kinesin activity is not sufficient to move the organelles outward as observed in fish melanophores (Fig. 1C, left vs. right panel). These findings suggest that amphibians rely on the functional cooperation between the microtubule and actin cytoskeletal systems for cell-wide redistribution of the melanosomes.

We previously provided a molecular rationale for these observations by studying the behavior of PKA-phosphorylated and dephosphorylated Rab27a/Mlph/MyoVa complexes from *Xenopus* frog and zebrafish at in vitro reconstituted microtubule-actin crossings (13). Consistent with the hypothesis that functional crosstalk between the microtubule and actin networks is required for the cell-wide redistribution of melanosomes in the frog model, the amphibian *Xl*Rab27a/*Xt*Mlph/*Xl*MyoVa complex displayed regulated switching at reconstituted microtubule-actin crossings in vitro, but not the homologous complex from fish (13). We eventually pinpointed this regulation to the FBD of the *Xt*Mlph adaptor protein in the amphibian transport complex (13). While the phosphorylation-state of *Xt*Mlph dictated the switching behavior of its *Xl*MyoVa at microtubule-actin crossings, up-regulation of switching onto microtubules in particular, it affected neither the speed nor the processivity of the motor on actin filaments (13). The latter suggests that the motor activity per se is not subject to phosphorylation-dependent regulation. Notably, this result was predicted by previous tracking of single melanosomes on the actin network in amphibian melanophores after removal of microtubules (14). In particular, theoretical modelling and simulations proposed that instead of regulating the motor activity, cells control the MyoVa-dependent switching of the organelles at actin-actin crossings to regulate the cell-wide transport (14). Upon increase of intracellular PKA activity, MyoVa was proposed to ignore the actin-actin crossings, increasing the displacement of melanosomes on the actin network (14). Conversely, upon decreased PKA activity, the motor was proposed to switch more frequently at actin-actin crossings, decreasing the overall displacement of melanosomes (14). In the simplest scenario, however, the homodimeric MyoVa motor that encounters an actin-actin crossing would statistically be expected to switch with an equal probability between the same filament types. Based on our previous demonstration of Mlph-regulated switching of the amphibian *Xl*Rab27a/*Xt*Mlph/*Xl*MyoVa complex at reconstituted microtubule-actin crossings (13), we hypothesized that the *Xt*Mlph adaptor might also regulate the switching behavior of its *Xl*MyoVa at actin-actin crossings. To directly address this question, and to test whether this presumed regulation entails any measurable impact on the overall distribution of melanosomes in melanophores, here we turned to in vitro reconstitution and in vivo assays, respectively.

## Results and Discussion

We previously demonstrated that the amphibian adaptor protein *Xt*Mlph from *Xenopus* frog interacts not only with actin filaments, but also with microtubules (13). The capability to interact with both filament types allowed *Xt*Mlph to regulate the switching probabilities of its *Xl*MyoVa at reconstituted microtubule-actin crossings (13). In stark contrast, *Dr*Mlph-a, *Dr*Mlph-b, and *Dr*Mlph-bX2 adaptor proteins from zebrafish failed to interact with microtubules (13). All three adaptor proteins consequently failed to regulate the switching probabilities of the *Dr*MyoVa motor at microtubule-actin intersections (13). Curiously, however, the *Dr*Mlph-bX2 acquired the capability to interact with actin filaments, as we demonstrated previously in our reconstitution assays (13). The fact that *Dr*Mlph-bX2, but not *Dr*Mlph-b, interacts with actin, is intriguing given the particularly high sequence identity between the two isoforms (95%, Fig. S1). Identification of such isoform- and species-specific properties provides us with an opportunity to identify the structural basis of differences in Mlph function and how this was altered during vertebrate evolution.

We therefore attempted to engineer an actin-binding capability into the *Dr*Mlph-b protein based on the sequence of the *Dr*Mlph-bX2 isoform (Fig. S1). It is well-established that a positively charged patch in the C-terminus of Mlph is important for actin binding (Fig. S2) (15-18). Indeed, a repetitive sequence of positively charged residues is present in the C-termini of both isoforms (highlighted in red in Fig. 2A), yet, only *Dr*Mlph-bX2 interacts with actin, and not the *Dr*Mlph-b isoform (Fig. 2B, left vs. middle panel). In contrast to *Dr*Mlph-b, the positively charged patch in *Dr*Mlph-bX2 is flanked with two prolines (highlighted in bold in Fig. 2A). We reasoned that these ‘rigid’ prolines might ‘loop out’ the positively charged patch and expose it for efficient actin binding. To test this hypothesis, we simply pasted this proline patch from *Dr*Mlph-bX2 into the *Dr*Mlph-b isoform (Fig. 2A). Remarkably, this replacement alone sufficed to impart a robust actin-binding capability onto the Mlph-b isoform (Fig. 2B, middle vs. right panel). Our findings thus suggest that a few strategic mutations are sufficient to impart filament-binding capability and thus allow a glimpse of how the Mlph protein may have started to acquire filament-binding capabilities over the course of evolution. However, whether the engineered actin-binding capability into the *Dr*Mlph-b protein can be ascribed solely to the exposed positively charged patch needs to be further studied in the future.

**Fig. 2:**
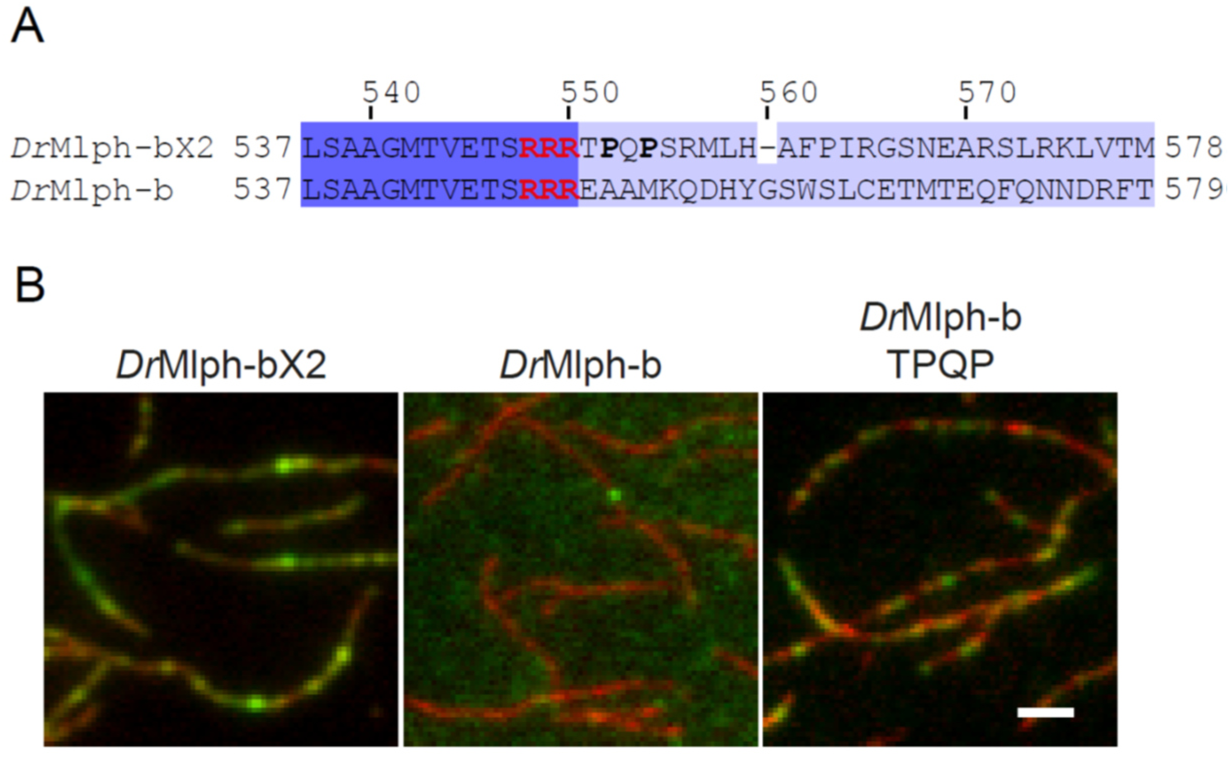
Engineering actin-binding capability into the *Dr*Mlph-b adaptor protein. (A) Sequence alignment of *Dr*Mlph-bX2 and *Dr*Mlph-b from zebrafish, respectively. The proposed actin-binding patch is highlighted in red in the sequences. (B) Decoration of surface-attached actin filaments (red) with fluorescently labeled wild-type and mutant *Dr*Rab27a/*Dr*Mlph complexes from zebrafish (green). Left panel shows the *Dr*Mlph-bX2 adaptor protein that is capable of interacting with actin filaments, which is not observed with *Dr*Mlph-b (middle panel). (Right panel) Introducing the prolines (highlighted in bold in (A)) into the *Dr*Mlph-b protein (*Dr*Mlph-b_EAA551-553TPQP) is sufficient to prompt actin binding. Scale bar: 2 µm.

We next asked whether the acquired capability of *Dr*Mlph-bX2 from zebrafish to bind actin filaments regulates the switching behavior of its *Dr*MyoVa at actin-actin crossings. To test this, we reconstituted actin-actin crossings in vitro and fluorescently labeled the *Dr*MyoVa from zebrafish as described previously (13). When the motor arrived at an actin-actin intersection, we simply scored whether it continued on its original filament or switched onto the crossing filament (Fig. 3, left panel). As statistically expected, the uncomplexed *Dr*MyoVa motor switched with equal probabilities at intersections that were reconstituted with the same filament type (Fig. 3A). To test our hypothesis whether the presence of the *Dr*Mlph-bX2 adaptor protein that acquired an actin-binding capability (Fig. 2B, left panel), gives rise to regulated switching at actin-actin crossings, we fluorescently labeled the *Dr*Rab27a subunit in the *Dr*Rab27a/*Dr*Mlph-bX2/*Dr*MyoVa complex. As a control, we additionally assembled the complex with the *Dr*Mlph-b adaptor that has not yet evolved the capability to interact with actin filaments. Given that the proposed switching is regulated by the increase and decrease of intracellular PKA activity (14), we phosphorylated and dephosphorylated the *Dr*Mlph-bX2 and the *Dr*Mlph-b adaptors in the respective complexes using PKA and antarctic phosphatase as described previously (13). Consistent with our previous observations at microtubule-actin crossings, neither complexes from zebrafish displayed regulated switching behavior and both switched with equal probabilities between the actin filaments (Fig. 3B and C, Movies 1-2). Taken together, our results indicate that acquisition of actin-binding by zebrafish Mlph isoforms is insufficient to allow them to undergo PKA-dependent track switching at actin-actin crossings.

**Fig. 3:**
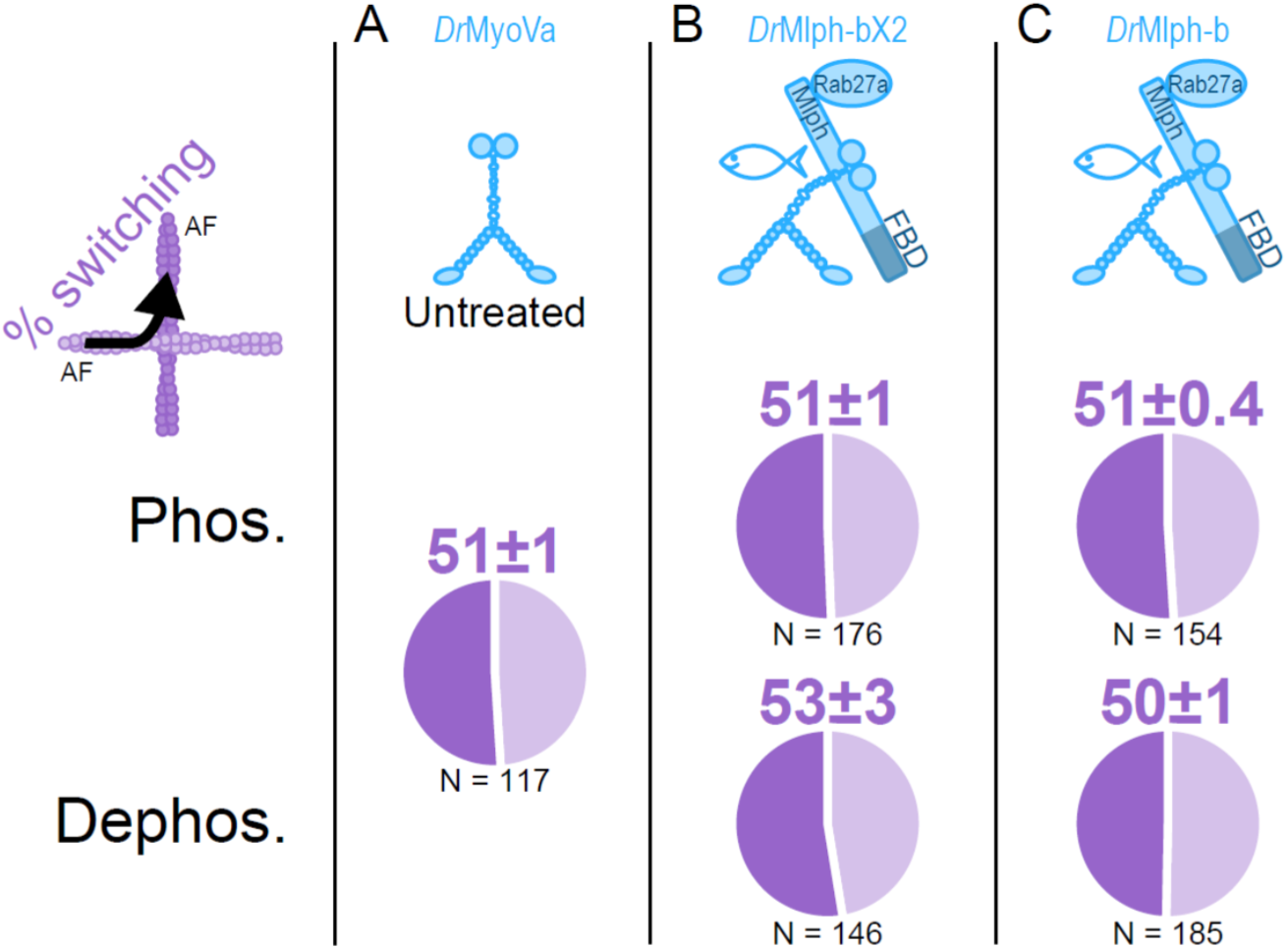
Mlph adaptors from zebrafish fail to regulate the switching of the *Dr*MyoVa motor at actin-actin crossings. (Top panel) Illustration of the *Dr*MyoVa motor and respective *Dr*Rab27a/*Dr*Mlph/*Dr*MyoVa complexes used in the study. The left column shows the switching direction and phosphorylation states (Phos. and Dephos.). Values (mean over experiments ± SD) above the pie charts represent the color-coded switching probabilities in percent. (A) Untreated *Dr*MyoVa motor switches with equal probabilities at actin-actin crossings as statistically expected. (B and C) *Dr*Mlph-bX2 that interacts with actin filaments, as well as *Dr*Mlph-b, both fail to regulate the switching of *Dr*MyoVa motor and the respective complexes recapitulate the switching behavior of *Dr*MyoVa shown in (A). N = number of events from 3 independent experiments. AF: actin filament.

We next turned to the amphibian model and asked whether the switching of the *Xl*Rab27a/*Xt*Mlph/*Xl*MyoVa complex from *Xenopus* frog is regulated by PKA at actin-actin crossings, as we demonstrated previously at microtubule-actin crossings (13). Strikingly, and in stark contrast to the zebrafish complex, about ∼2/3 of the PKA-phosphorylated *Xenopus* complexes ignored the actin-actin crossings and continued on the original filament (Fig. 4A, top panel, Phos.). Dephosphorylation completely reversed this behavior and ∼2/3 of the complexes switched at the respective intersections (Fig. 4A, bottom panel, Dephos.). Further consistent with our initial hypothesis of Mlph-mediated regulation of switching behavior, removal of the C-terminal FBD in *Xt*Mlph abolished this regulation and the *Xl*MyoVa motor switched with equal probabilities between the actin filaments (Fig. 4B, top vs. bottom panel). The next non-trivial task will be to unmask the structural determinants of the interaction between the *Xt*Mlph and actin filaments to fully understand how the adaptor protein controls the switching propensity of its *Xl*MyoVa motor at actin-actin intersections.

**Fig. 4:**
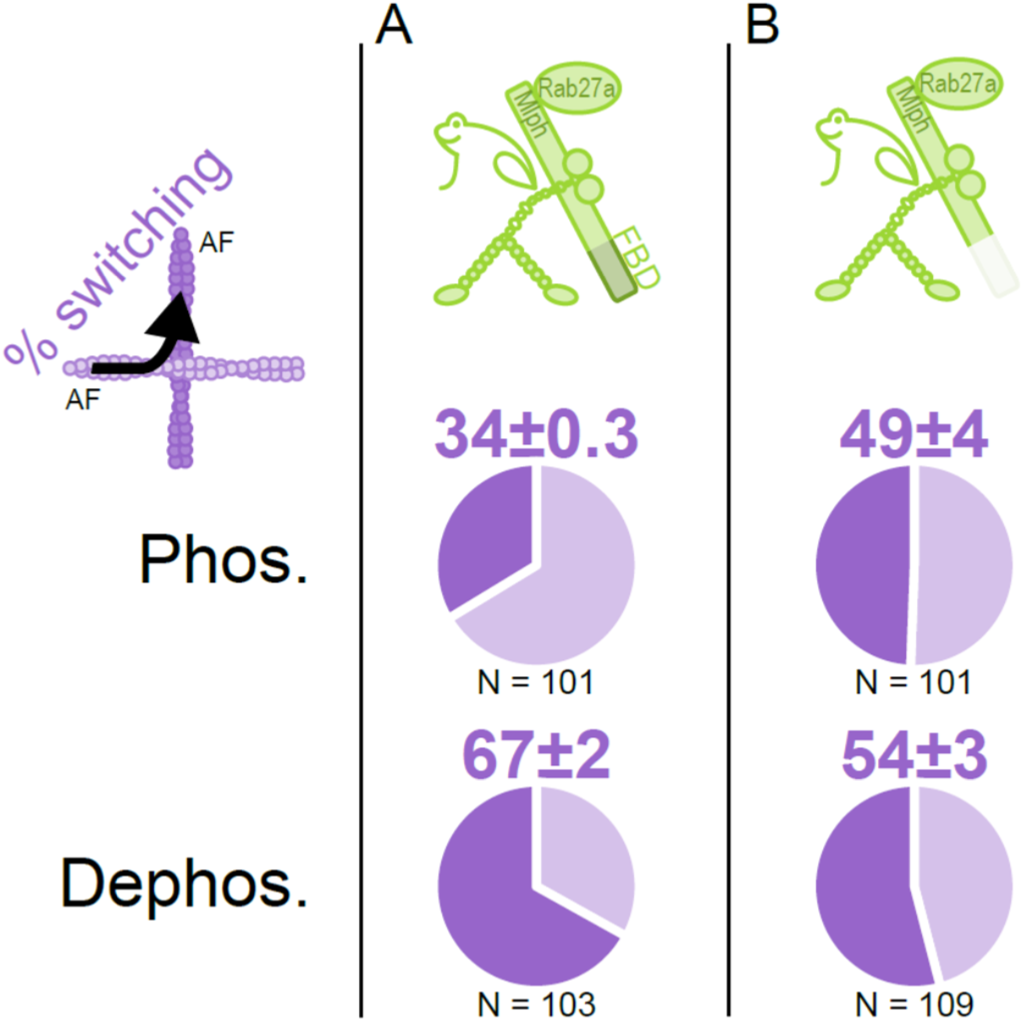
FBD of the amphibian *Xt*Mlph regulates the switching behavior of the *Xl*MyoVa motor at actin-actin crossings. (Top panel) Illustration of the *Xl*Rab27a/*Xt*Mlph/*Xl*MyoVa complexes used in the study. The left column shows the switching direction and phosphorylation states (Phos. and Dephos.). Values (mean over experiments ± SD) above the pie charts represent the color-coded switching probabilities in percent. (A) (Top vs. bottom panel) Only ∼1/3 of the PKA-phosphorylated *Xl*Rab27a/*Xt*Mlph/*Xl*MyoVa complexes switch at actin-actin crossings (top), and ∼2/3 ignore the intersections and continue on their original filament. (Bottom) About ∼2/3 of the dephosphorylated complexes switch onto the crossing filament, reversing the effect of PKA-dependent phosphorylation. (B) (Top vs. bottom panel) Removal of the FBD abolishes the regulated switching of the *Xl*MyoVa motor and the complexes switch with equal probabilities between the actin filaments. N = number of events from 3 independent experiments. AF: actin filament.

Lastly, we turned to *Xenopus* melanophores and asked whether this regulated switching of MyoVa at actin crossings, as proposed by previous simulations (14), and here directly demonstrated in in vitro reconstitution experiments (Fig. 4), affects the regulated redistribution of melanosomes on the actin network. To this end, we tested the effect of PKA-elevating and -lowering agents (MSH and melatonin) on melanosome distribution in microtubule-depleted *Xenopus* melanophores. We first confirmed the results of previous work (11, 19-26) showing that MSH and melatonin cause melanosome dispersion and clustering, respectively (Fig. 5A and B, row 1 vs. 2), and that clustering requires intact microtubules as (Fig. 5A and B, compare row 1 vs. 3). This indicates that melanosome dispersion seen in cells in row 3 reflects MyoVa/actin-dependent melanosome transport. Consistent with this notion, disruption of the actin network by cytochalasin-D treatment suppressed this dispersion and melanosomes remained clustered (Fig. 5A and B, row 3 vs. 4).

**Fig. 5:**
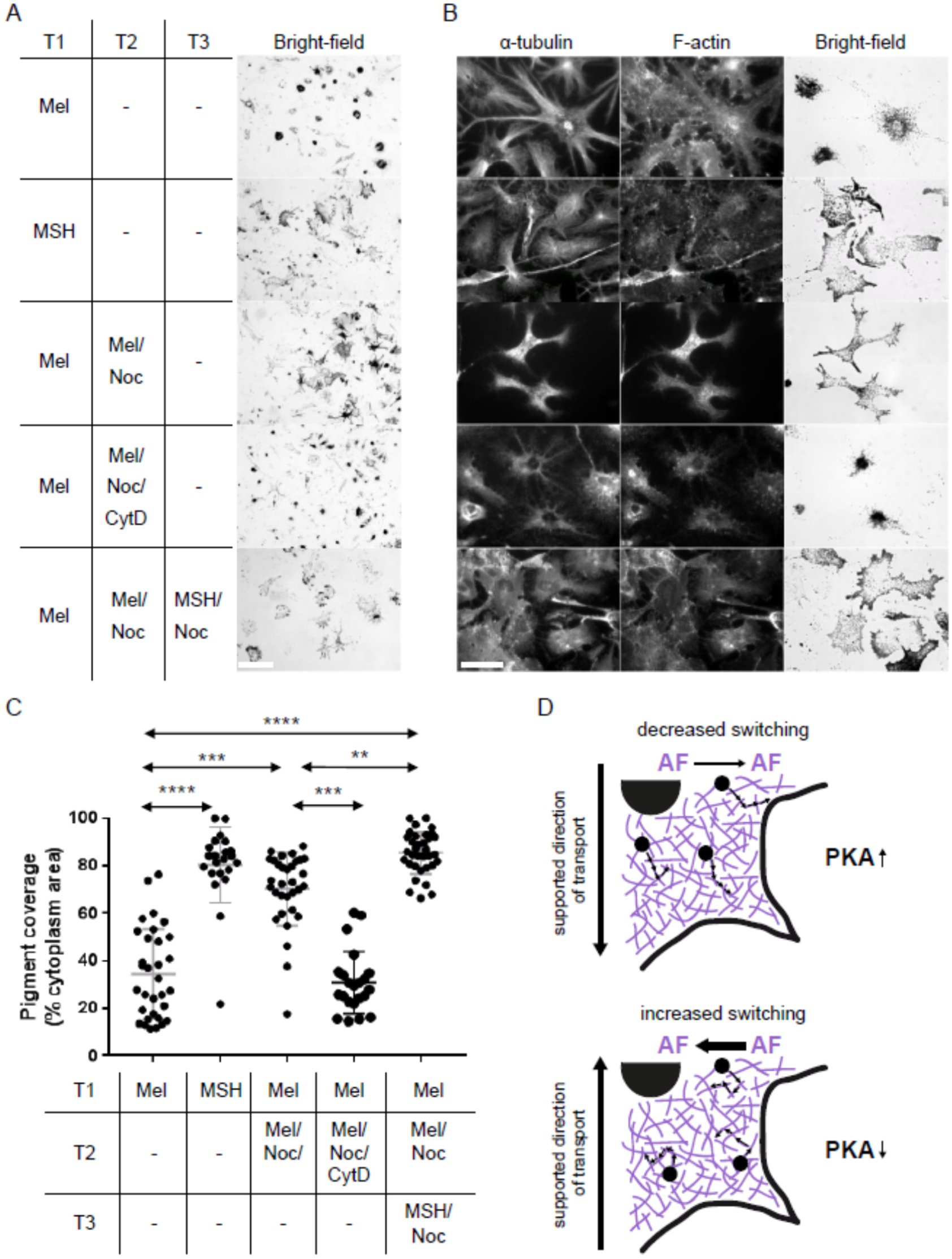
MSH significantly enhances melanosome dispersion in microtubule-depleted *Xenopus* melanophores *in vivo*. Melanophores were plated onto glass cover-slips, serum-starved overnight, incubated with the indicated combination of melatonin/MSH, and nocodazole/cytochalasin D, respectively, fixed after the final incubation and prepared for immunofluorescence microscopy (see Materials and Methods for details). Each incubation was for 1 hour at 27 °C or 4 °C for conditions including nocodazole. **(A)** Left panel shows the regime of incubations and the right panel shows the low magnification bright-field images of distribution of melanosomes in populations of cells in each condition, respectively. **(B)** High magnification bright-field and fluorescence images showing the distribution of melanosomes, microtubules and actin filaments in cells. **(C)** A scatter-plot showing the distribution of melanosomes in populations of melanophores. N = 32 (melatonin and melatonin/nocodazole), 22 (MSH), 22 (melatonin/nocodazole/cytochalasin D), and 34 (MSH/nocodazole). Data was analyzed using one-way ANOVA followed by Kruskal-Wallis post-test ** = p<0.01, *** = p<0.001 and **** = p<0.0001. No other population comparisons yielded significant differences. Bars, 100 µm (A) and 40 µm (B). **(D)** Illustration of the switching regulated displacement of melanosomes as a response to intracellular PKA activity in melanophores. (Top panel) Increase of PKA activity decreases the switching probability of MyoVa and enhances the displacement of the melanosomes for dispersion. (Bottom panel) Decrease of PKA activity (i.e. transport towards the cell center for melanosome clustering) increases the switching probability and decreases the overall displacement of the organelles and contributes to aggregation. AF: actin filament.

The MyoVa-mediated dispersal seen in row 3 results from the decreased PKA and increased phosphatase activities, respectively. According to our experimental findings (Fig. 4) and previous simulations (14), this cell state corresponds to the dephosphorylated Rab27a/Mlph/MyoVa complex with increased switching probability at actin-actin crossings (Fig. 4A, bottom panel, Dephos.). As such, the degree of dispersion should be decreased due to the frequent switching of the MyoVa motor (14). If so, solely increasing the PKA activity, thus decreasing the MyoVa switching (Fig. 4A, top panel, Phos.), is expected to further increase the degree of dispersal seen in row 3. Remarkably, MSH-dependent increase of PKA activity significantly enhanced melanosome dispersion (Fig. 5A and B, row 3 vs. 5), indicating that reduced switching on the actin network enhances MyoVa/actin-driven melanosomes dispersion. Based on our reconstitution studies and consistent with previous predictions, we conclude that observed differences with respect to the displacement of melanosomes result from the Mlph-regulated switching of MyoVa at actin-actin crossings, and not from the regulation of the motor’s activity.

Together with our previous findings, we propose that the regulation of MyoVa switching at microtubule-actin and actin-actin crossings both contribute to the correct positioning of melanosomes on the cytoskeleton that we now collectively trace back to the FBD of the Mlph adaptor protein (Fig. 5D). This FBD-mediated control of switching at cytoskeletal crossings observed with the amphibian *Xl*Rab27a/*Xt*Mlph/*Xl*MyoVa complex (Fig. 4) (13) is, however, completely absent in the zebrafish complex (Fig. 3). The lack of response of the zebrafish *Dr*Rab27a/*Dr*Mlph/*Dr*MyoVa complex to PKA also at microtubule-actin crossings (13) together suggests that the MyoVa/actin system is not yet under the control of the PKA-dependent regulation in the fish model. Previous in vivo experiments in fact delivered first indications that it is the microtubule, and not the actin network system, that is subject to regulation in fish melanophores (27). Consistently, and in stark contrast to amphibians, the microtubule-based transport system alone is sufficient for hyperdispersion of melanosomes (Fig. 1C, left panel), and as such, for the voluntary color change in the fish model. Our dissection of the amphibian *Xl*Rab27a/*Xt*Mlph/*Xl*MyoVa complex, on the other hand, suggests that the evolution of the Mlph adaptor protein, its C-terminal FBD in particular, eventually established a functional response to the changes of intracellular PKA activity. FBD specifically controls the switching behavior of the MyoVa motor not only at microtubule-actin (13) but also at actin-actin crossings that we now reconstituted in our functional assays (Fig. 4). Importantly, both types of regulated switching on the cytoskeleton support the correct overall direction of transport (Fig. 5D). Together with previous findings, data presented here consistently point to a direct link between the cAMP-PKA regulatory pathway and the myosin-driven transport complex. Notably, however, this functional link does not ‘up- and down’ regulate the activity of the motor, but acts as a global navigation system throughout the entire cytoskeletal lattice as a response to the changes of intracellular PKA activity.

## Data availability

All data that support the findings of this study are available within the paper and its SI Appendix.

## Acknowledgements

We thank Vladimir I. Gelfand (Northwestern University, Chicago, U.S.A.) for his generous gift of melanophore culture, and Vladimir I. Gelfand (Northwestern University, Chicago, U.S.A.) and Minjong Park (University of California, San Francisco, U.S.A.) for the melanophilin plasmids. We are grateful to Günther Woehlke (Technische Universität München, Munich, Germany), and all members of the Ökten group for fruitful discussions on the project. This work was funded by the Deutsche Forschungsgemeinschaft (DFG, German Research Foundation) – SFB-863 – Project ID 111166240. Z.Ö. acknowledges a starting grant from the European Research Council (GA no. 335623). A.N.H. acknowledges support from MRC New Investigator Award G1100063 and Wellcome Trust Institutional Strategic Support Fund award 204843/Z/16/Z.

## Author Contributions

A.O., P.S., D.Z., A.N.H. and Z.Ö. planned the experiments. A.O., P.S., D.Z., E.B., and A.N.H. performed experiments. A.O., P.S., D.Z. and A.N.H. analyzed data. A.O., A.N.H., and Z.Ö. wrote the manuscript.

## Materials and Methods

### Reagents

All reagents were the highest purity available and were obtained from Sigma-Aldrich (Munich, Germany) or Carl Roth (Karlsruhe, Germany), unless mentioned otherwise.

### DNA constructs

All constructs were cloned into the vector pFastBacI for subsequent expression in the baculovirus system (Life Technologies, Darmstadt, Germany). A number of constructs were codon-optimized for expression in insect cells and synthesized commercially (GenScript, Piscataway, U.S.A.). *Dr*Rab27a, *Dr*Mlph-b, *Dr*Mlph-bX2, *Dr*MyoVa, *Xl*Rab27a, *Xt*Mlph, *Xt*MlphΔFBD, and *Xl*MyoVa were described earlier (13, 28). *Dr*Mlph-b-TPQP was synthesized with residues 551 to 554 changed to TPQP.

### Protein expression, purification, fluorescent labeling, and dephosphorylation/phosphorylation and reconstitution of tripartite complexes

All procedures were performed as described in (13, 28).

### Sequence alignment

*Mm*Mlph (NP_443748.2), *Hs*Mlph (NP_077006.1), *Fd*Mlph (XP_010624436.1), *Cl*Mlph (NP_001096689.2), *Oa*Mlph (NP_001139743.1), *Xt*Mlph (NP_001120194.1), *Dr*Mlph-a (XP_005168769.1), *Dr*Mlph-b (XP_021334596.1), and *Dr*Mlph-bX2 (XP_021334597.1) were aligned using Jalview 2.10.5 software (29) and the web service T-Coffee (30) using default parameters except gap penalty (set to 0).

### Reconstitution assay with tripartite complexes on microtubules and actin filaments

Biotinylated Atto488-labeled microtubules and biotinylated Atto565-labeled actin filaments were attached to the surface of a flow chamber. Alexa Fluor 647-labeled dephosphorylated or phosphorylated Rab27a/Mlph/MyoVa complexes were flowed into these chambers and reconstitution assays were carried out with 4 mM ATP as described previously (28). Movies were acquired at room temperature with the microscope setup described earlier (28). To analyze the switching behavior of the MyoVa transport complexes on actin networks, only complexes in close proximity to an actin-actin crossing were taken into account. The number of complexes that switched filament at a crossing or passed the crossing without changing filament was manually counted. Complexes that stopped their movement on a crossing were not considered as events. The switching probabilities (i.e. number of complexes that switched at actin-actin crossings divided by total number of complexes that went across crossings in %) were determined for each complex (Dephos. And Phos.). We obtained individual switching probabilities from three independent measurements. Given switching probabilities were calculated as mean values of these individually determined switching probabilities and standard deviations were calculated using

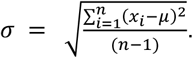

### Melanosome distribution assay

For studies of melanosome distribution, melanophores were plated at a density of 1×104 cells/ml in growth medium into wells of a 24-well cell culture plate that were pre-loaded with uncoated 13 mm glass coverslips. After 24 hours, the medium was replaced with serum-free medium. Sixteen hours later this was replaced with serum-free medium supplemented with melatonin (100 nM) or MSH (10 nM) in the presence or absence of nocodazole (10 µM) or cytochalasin D (5 µg/ml), respectively. All incubations were for 1 hour at 27°C or 4°C for nocodazole conditions, respectively. Cells were then fixed and processed for immunofluorescence as described previously (31, 32). Microtubules were detected using mouse-anti-tubulin (DM1a; Calbiochem cp06) followed by Alexa488 conjugated goat anti-mouse secondary antibodies (Invitrogen A-11001) diluted at 1:100 and 1:500, respectively. F-actin was detected as previously described using Texas-red-conjugated phalloidin. Images showing melanosome, microtubule and F-actin distribution in melanophores were captured using an Axiovert S-100 microscope as described previously (33). Analysis of intracellular melanosome distribution was performed as described previously (16). Briefly, the total and melanosome filled area for individual cells were determined manually using ImageJ and pigment coverage was determined by calculating the percentage of total cell area occupied by melanosomes.

## References

1. A. N. Hume, M. C. Seabra, Melanosomes on the move: a model to understand organelle dynamics. Biochem Soc Trans 39, 1191–1196 (2011).

2. A. A. Nascimento, J. T. Roland, V. I. Gelfand, Pigment cells: a model for the study of organelle transport. Annu Rev Cell Dev Biol 19, 469–491 (2003).

3. H. Nilsson Skold, S. Aspengren, M. Wallin, Rapid color change in fish and amphibians - function, regulation, and emerging applications. Pigment Cell Melanoma Res 26, 29–38 (2013).

4. V. I. Rodionov, F. K. Gyoeva, V. I. Gelfand, Kinesin is responsible for centrifugal movement of pigment granules in melanophores. Proc Natl Acad Sci U S A 88, 4956–4960 (1991).

5. S. L. Rogers, V. I. Gelfand, Myosin cooperates with microtubule motors during organelle transport in melanophores. Curr Biol 8, 161–164 (1998).

6. S. L. Rogers, I. S. Tint, P. C. Fanapour, V. I. Gelfand, Regulated bidirectional motility of melanophore pigment granules along microtubules in vitro. Proc Natl Acad Sci U S A 94, 3720–3725 (1997).

7. M. C. Tuma, A. Zill, N. Le Bot, I. Vernos, V. Gelfand, Heterotrimeric kinesin II is the microtubule motor protein responsible for pigment dispersion in Xenopus melanophores. J Cell Biol 143, 1547–1558 (1998).

8. A. Daniolos, A. B. Lerner, M. R. Lerner, Action of light on frog pigment cells in culture. Pigment Cell Res 3, 38–43 (1990).

9. T. L. Eriksson et al., Panax ginseng induces anterograde transport of pigment organelles in Xenopus melanophores. J Ethnopharmacol 119, 17–23 (2008).

10. M. C. Isoldi, I. Provencio, A. M. Castrucci, Light modulates the melanophore response to alpha-MSH in Xenopus laevis: an analysis of the signal transduction crosstalk mechanisms involved. Gen Comp Endocrinol 165, 104–110 (2010).

11. A. R. Reilein, I. S. Tint, N. I. Peunova, G. N. Enikolopov, V. I. Gelfand, Regulation of organelle movement in melanophores by protein kinase A (PKA), protein kinase C (PKC), and protein phosphatase 2A (PP2A). J Cell Biol 142, 803–813 (1998).

12. V. I. Rodionov, A. J. Hope, T. M. Svitkina, G. G. Borisy, Functional coordination of microtubule-based and actin-based motility in melanophores. Curr Biol 8, 165–168 (1998).

13. A. Oberhofer et al., Molecular underpinnings of cytoskeletal cross-talk. Proc Natl Acad Sci U S A 117, 3944–3952 (2020).

14. J. Snider et al., Intracellular actin-based transport: How far you go depends on how often you switch. P Natl Acad Sci USA 101, 13204–13209 (2004).

15. M. Fukuda, T. S. Kuroda, Slac2-c (synaptotagmin-like protein homologue lacking C2 domains-c), a novel linker protein that interacts with Rab27, myosin Va/VIIa, and actin. J Biol Chem 277, 43096–43103 (2002).

16. A. N. Hume, A. K. Tarafder, J. S. Ramalho, E. V. Sviderskaya, M. C. Seabra, A coiledcoil domain of melanophilin is essential for Myosin Va recruitment and melanosome transport in melanocytes. Mol Biol Cell 17, 4720–4735 (2006).

17. T. S. Kuroda, H. Ariga, M. Fukuda, The actin-binding domain of Slac2-a/melanophilin is required for melanosome distribution in melanocytes. Mol Cell Biol 23, 5245–5255 (2003).

18. M. Sckolnick, E. B. Krementsova, D. M. Warshaw, K. M. Trybus, More than just a cargo adapter, melanophilin prolongs and slows processive runs of myosin Va. J Biol Chem 288, 29313–29322 (2013).

19. K. Abe et al., Role of cyclic AMP in mediating the effects of MSH, norepinephrine, and melatonin on frog skin color. Endocrinology 85, 674–682 (1969).

20. P. N. de Graan, A. N. Eberle, Irreversible stimulation of Xenopus melanophores by photoaffinity labelling with p-azidophenylalanine13-alpha-melanotropin. FEBS Lett 116, 111–115 (1980).

21. B. Magun, Two actions of cyclic AMP on melanosome movement in frog skin. Dissection by cytochalasin B. J Cell Biol 57, 845–858 (1973).

22. M. N. Potenza, M. R. Lerner, A rapid quantitative bioassay for evaluating the effects of ligands upon receptors that modulate cAMP levels in a melanophore cell line. Pigment Cell Res 5, 372–378 (1992).

23. D. Sugden, S. J. Rowe, Protein kinase C activation antagonizes melatonin-induced pigment aggregation in Xenopus laevis melanophores. J Cell Biol 119, 1515–1521 (1992).

24. T. Ebisawa, S. Karne, M. R. Lerner, S. M. Reppert, Expression cloning of a high-affinity melatonin receptor from Xenopus dermal melanophores. Proc Natl Acad Sci U S A 91, 6133–6137 (1994).

25. A. B. Lerner, J. D. Case, Y. Takahashi, Isolation of melatonin and 5-methoxyindole-3-acetic acid from bovine pineal glands. J Biol Chem 235, 1992–1997 (1960).

26. B. H. White, R. D. Sekura, M. D. Rollag, Pertussis toxin blocks melatonin-induced pigment aggregation in Xenopus dermal melanophores. J Comp Physiol B 157, 153–159 (1987).

27. B. M. Slepchenko, I. Semenova, I. Zaliapin, V. Rodionov, Switching of membrane organelles between cytoskeletal transport systems is determined by regulation of the microtubule-based transport. J Cell Biol 179, 635–641 (2007).

28. A. Oberhofer et al., Myosin Va’s adaptor protein melanophilin enforces track selection on the microtubule and actin networks in vitro. P Natl Acad Sci USA 114, E4714–E4723 (2017).

29. A. M. Waterhouse, J. B. Procter, D. M. Martin, M. Clamp, G. J. Barton, Jalview Version 2--a multiple sequence alignment editor and analysis workbench. Bioinformatics 25, 1189–1191 (2009).

30. C. Notredame, D. G. Higgins, J. Heringa, T-Coffee: A novel method for fast and accurate multiple sequence alignment. J Mol Biol 302, 205–217 (2000).

31. A. N. Hume, D. S. Ushakov, A. K. Tarafder, M. A. Ferenczi, M. C. Seabra, Rab27a and MyoVa are the primary Mlph interactors regulating melanosome transport in melanocytes. J Cell Sci 120, 3111–3122 (2007).

32. R. D. Evans et al., Myosin-Va and dynamic actin oppose microtubules to drive longrange organelle transport. Curr Biol 24, 1743–1750 (2014).

33. C. L. Robinson, R. D. Evans, D. A. Briggs, J. S. Ramalho, A. N. Hume, Inefficient recruitment of kinesin-1 to melanosomes precludes it from facilitating their transport. J Cell Sci 130, 2056–2065 (2017).

