## Supplementary material for "How to navigate the myosin-V motor through the actin network": SI Appendix

### Supplementary Figures

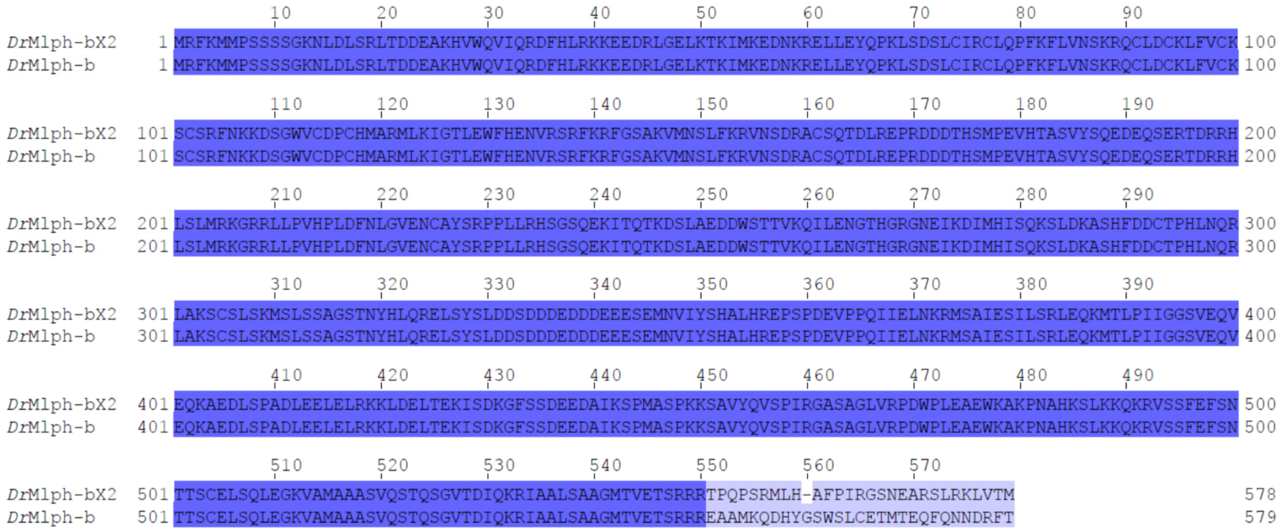


**Fig. S1: Sequence alignment of the isoforms *Dr*Mlph-b and *Dr*Mlph-bX2.** The isoforms are identical except for 30 amino acids at the C-terminus. Both proteins contain the positively charged patch (R548/549/550). *Dr*Mlph-bX2 additionally possesses two proline residues in close proximity of the arginine repeats that might expose this positively charged patch and enable actin binding.

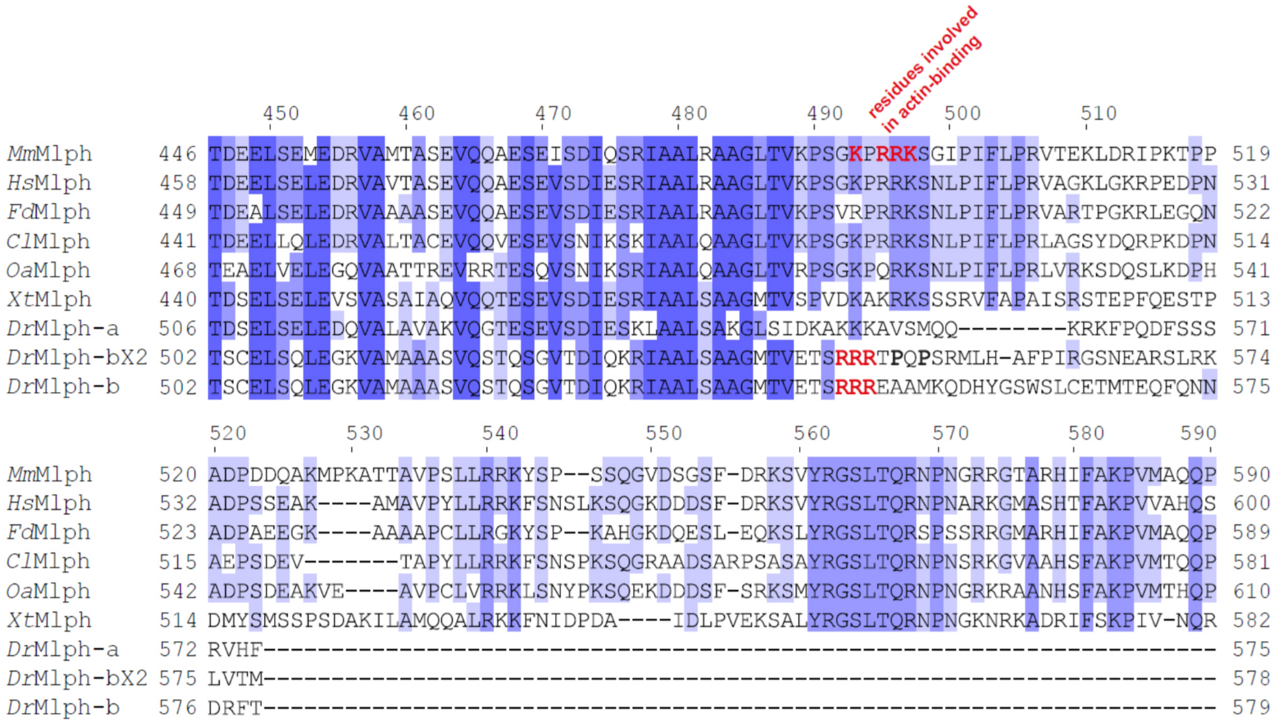


**Fig. S2:** Sequence alignment of the FBDs from mouse (*Mm*), human (*Hs*), mole-rat (*Fd*), dog (*Cl*), sheep (*Oa*), *Xenopus* (*Xt*), and zebrafish (*Dr*); numbers above correspond to positions in the mouse Mlph. Positions for each Mlph are given at left and right of alignment. Residues (K493, R495, R496, K497) involved in actin binding in *Mm*Mlph were identified previously (1-4) and are indicated in red. Patch of positively charged amino acids in *Dr*Mlph-b and *Dr*MlphbX2 are shown in red. Proline residues that C-terminally flank the putative actin-binding site in DrMlph-bX2 are highlighted in bold.

### Supplementary Movie Legends

**Movie S1: Example of a MyoVa transport complex from zebrafish that switches the actin filament at an actin-actin crossing in vitro.** The tripartite complex labeled via the Rab27a subunit is shown in green and actin filaments in red and microtubules in blue. Movie is displayed at 1.21x real time. Scale bar: 2 µm.

**Movie S2: Example of a MyoVa transport complex from zebrafish that passes an actin-actin crossing without switching filament and continues on the same filament in vitro.** Colors as in Movie S1. Movie is displayed at 1.21x real time. Scale bar: 2 µm.

**Movie S3: Example of a MyoVa transport complex from *Xenopus* frog that switches the actin filament at an actin-actin crossing in vitro.** Colors as in Movie S1. Movie is displayed at 1.21x speed. Scale bar: 2 µm.

**Movie S4: Example of a MyoVa transport complex from zebrafish that passes an actin-actin crossing without switching filament and continues on the same filament in vitro.** Colors as in Movie S1. Movie is displayed at 1.21x real time. Scale bar: 2 µm.

### References

1. M. Fukuda, T. S. Kuroda, Slac2-c (synaptotagmin-like protein homologue lacking C2 domains-c), a novel linker protein that interacts with Rab27, myosin Va/VIIa, and actin. *J Biol Chem* **277**, 43096-43103 (2002).

2. A. N. Hume, A. K. Tarafder, J. S. Ramalho, E. V. Sviderskaya, M. C. Seabra, A coiled-coil domain of melanophilin is essential for Myosin Va recruitment and melanosome transport in melanocytes. *Mol Biol Cell* **17**, 4720-4735 (2006).

3. T. S. Kuroda, H. Ariga, M. Fukuda, The actin-binding domain of Slac2-a/melanophilin is required for melanosome distribution in melanocytes. *Mol Cell Biol* **23**, 5245-5255 (2003).

4. M. Sckolnick, E. B. Krementsova, D. M. Warshaw, K. M. Trybus, More than just a cargo adapter, melanophilin prolongs and slows processive runs of myosin Va. *J Biol Chem* **288**, 29313-29322 (2013).
